# Multiple parasitism promotes facultative host acceptance of cuckoo eggs and rejection of cuckoo chicks

**DOI:** 10.1101/2021.08.18.456772

**Authors:** Hee-Jin Noh, Ros Gloag, Naomi E. Langmore

**Author notes:** **Correspondence** Hee-Jin Noh.

## Abstract

Many hosts of brood parasitic cuckoos reject foreign eggs from the nest. Yet where nests commonly receive more than one cuckoo egg, hosts might benefit by instead accepting parasite eggs. This is because cuckoos remove an egg from the nest before adding their own, and keeping cuckoo eggs in the nest reduces the odds that further host eggs are removed by subsequent cuckoos. This ‘clutch dilution effect’ has been proposed as a precondition for the evolution of cuckoo nestling eviction by hosts, but no previous studies have tested this in a host that rejects cuckoo nestlings. We tested the clutch dilution hypothesis in large-billed gerygones (*Gerygone magnirostris*), which are multiply parasitized by little bronze-cuckoos (*Chalcites minutillus*). Gerygones evict cuckoo nestlings but accept cuckoo eggs. Consistent with multiple parasitism favouring egg acceptance, we found gerygone egg survival was higher under scenarios of cuckoo egg acceptance than rejection. Yet gerygones were also flexible in their egg acceptance, with 35% abandoning cuckoo-egg-only clutches. This novel demonstration of adaptive clutch dilution suggests that multiple parasitism can favour a facultative response to brood parasite eggs, whereby hosts accept or reject parasite eggs depending on clutch composition.

## Introduction

The interactions between brood parasites and their hosts are classic examples of coevolutionary arms races; hosts evolve defenses against parasites, which then select for counter-adaptations in the parasite (Davies, 2000; Davies, 2011; Rothstein, 1990). For example, many hosts of brood parasitic birds have evolved the ability to recognize and reject parasite eggs (Davies and Brooke, 1989, Spottiswoode and Stevens, 2010), in turn selecting for mimicry of host eggs in brood parasites (Stoddard and Stevens, 2010, 2011; Attard et al., 2017). Similarly, other hosts have evolved the ability to recognize and reject parasite nestlings (Langmore et al., 2003, Sato et al., 2010; Tokue and Ueda, 2010), which has selected for mimicry of host nestlings by brood parasites (Langmore et al., 2011; Noh et al., 2018; Attisano et al., 2018). However, our understanding of these coevolutionary interactions is challenged when some hosts fail to evolve defences against their parasite, despite high costs of parasitism.

One puzzle is why egg rejection and chick rejection are never observed in the same host. Chick rejection is rare, but is common among hosts of three species of bronze-cuckoos *Chalcites* spp. (Langmore et al., 2003; Sato et al., 2010; Tokue and Ueda, 2010; Sato et al., 2015). One explanation for this is that egg rejection is always the better strategy for hosts, and that effective egg rejection then reduces selection for subsequent recognition and removal of chicks. Chick rejection, therefore, should evolve *only* in systems where egg ejection has a prohibitively high cost (Britton et al., 2007; Grim, 2006; Planqué et al., 2002) or is constrained by the nest environment (Langmore et al., 2009). While several potential costs of egg ejection have been identified (e.g. recognition error, Marchant, 1972; Brooker et al., 1990; Langmore et al., 2005; accidental damage to host eggs; Antonov et al., 2006; Lorenzana and Sealy, 2001), it has yet to be demonstrated that egg ejection is costly in any known chick-ejecting hosts.

Here, we test the idea that a cost of rejecting parasitic eggs is the precondition for the evolution of cuckoo chick ejection by hosts (Sato et al., 2010), via a field study of Australia’s large-billed gerygone (*Gerygone magnirostris*), a chick-ejecting host of the little bronze-cuckoo (*Chalcites minutillus*). When individual female cuckoos overlap in their use of hosts, hosts may regularly receive two or more parasite eggs in the nest. Under these circumstances, acceptance of the parasite eggs can yield a better pay-off for the host than egg eviction due to a ‘clutch dilution effect’ (Sato et al., 2010). This is because the parasite typically removes or destroys one or more eggs in the nest prior to laying her own egg, such that later parasites sometimes remove the eggs laid by earlier ones, rather than removing the host’s own eggs (Davies and Brooke, 1988), thereby reducing the risk of host egg loss during the egg stage. The benefits of clutch dilution have been demonstrated in a host of a non-evicting parasite (the shiny cowbird, *Molothrus bonariensis*) where host young are reared alongside the parasite young (Gloag et al., 2012). In this case, the benefit of retaining parasite eggs is presumably sufficient to offset the cost of rearing parasite chicks (Gloag et al., 2012). By contrast, for large-billed gerygones and other species exploited by cuckoos that evicts host chicks soon after hatching, host parents can enjoy a clutch dilution effect of tolerating parasite eggs only if they instead defer parasite rejection to the chick stage (Sato et al., 2010).

Large-billed gerygones appear to provide a good fit for the conditions of the clutch dilution hypothesis. They do not reject cuckoo eggs, but can rescue a parasitized brood by recognizing and evicting cuckoo chicks from the nest soon after hatching (Noh et al., 2018; Sato et al., 2009). Large-billed gerygones have a small clutch size of 2-3 eggs and their nests are regularly exploited by multiple female cuckoos (Gloag et al., 2014). Also, little bronze-cuckoos nearly always remove one egg at the time of parasitism (83.3% parasitism events; Gloag et al., 2014). Little bronze-cuckoos lay a dark olive or brown coloured egg, which is cryptic inside the dome-shaped nests of large-billed gerygones and quite distinct from the speckled white eggs of the host (Langmore et al., 2009). Cuckoo egg crypsis may lead second-to-arrive cuckoos to bias their egg removal toward host eggs (Gloag et al., 2014), but egg acceptance will still bring a net benefit to gerygones provided cuckoos sometimes remove previously-laid cuckoo eggs.

We assessed whether retaining cuckoo eggs in the nest increases gerygone egg survivorship, relative to rejecting cuckoo eggs, by comparing the number of surviving gerygone eggs in a clutch after second parasitism events. In addition, we extend the clutch dilution hypothesis by considering an additional scenario, in which high rates of multiple parasitism result in an increased probability of nests containing only cuckoo eggs. For example, among 2-egg gerygone clutches, parasitism by two cuckoos will sometimes result in a clutch with two cuckoo eggs and no gerygone eggs. In such cases, accepting cuckoo eggs would no longer be beneficial, and hosts would benefit by switching to alternative defences, such as nest abandonment or clutch rejection (De Mársico et al., 2013; Langmore et al., 2003). This type of flexible strategy would depend on hosts being able to detect that their nest contains either only parasite eggs, or no host eggs.

## Methods

We conducted fieldwork along creek lines in the Cairns region (16°55′ S, 145°46′ E), Queensland, Australia. We searched for large-billed gerygone nests during the breeding season (Aug-Dec, 2016-2018), and monitored the incidence and intensity of parasitism.

Large-billed gerygones lay one egg every second day to produce a typical clutch of 2-3 eggs (mean: 3±0.09 eggs, range: 1-5, n = 100; Noh et al., 2018). During egg-laying, we visited the nests every day and marked eggs to identify them and to check for parasitism, egg rejection, and nest abandonment. We then continued monitoring the nests at intervals of four- or five-days during the incubation and nestling stages.

To assess whether the presence of cuckoo eggs in the clutch decreases the risk of gerygone egg loss due to subsequent parasitism, we selected nests with three clutch size and assessed the number of gerygone eggs remaining in nests after second parasitism events for two categories of multiply parasitized clutches: (A) “Multiple parasitism egg accepters” (n=13); nests that had already lost one gerygone egg to cuckoo parasitism (i.e. clutches of two gerygone eggs plus a cuckoo egg) at the time of a second parasitism event. (B) “Virtual egg rejecters” (n=14); nests that contained two gerygone eggs at the time of the second parasitism event. This group represented the outcomes for a parasitized three-gerygone-egg clutch in which the first cuckoo egg had been rejected rather than accepted (Fig. 1-a). Higher gerygone egg survival in nests of our “egg acceptor” group than our “virtual egg rejecter” group would indicate that cuckoo eggs in the nest can protect host egg survival, and thus support the clutch dilution hypothesis for parasite egg acceptance in this host. For comparison, we also recorded the number of gerygone eggs remaining in unparasitised nests ((C) “Unparasitised” (n=26); all unparasitized nests with a clutch size of three) and in nests parasitized just once ((D) “Single parasitism” (n=28); nests that contained three gerygone eggs at the time of parasitism by a single cuckoo) (Fig. 1-a).

**Figure 1.**
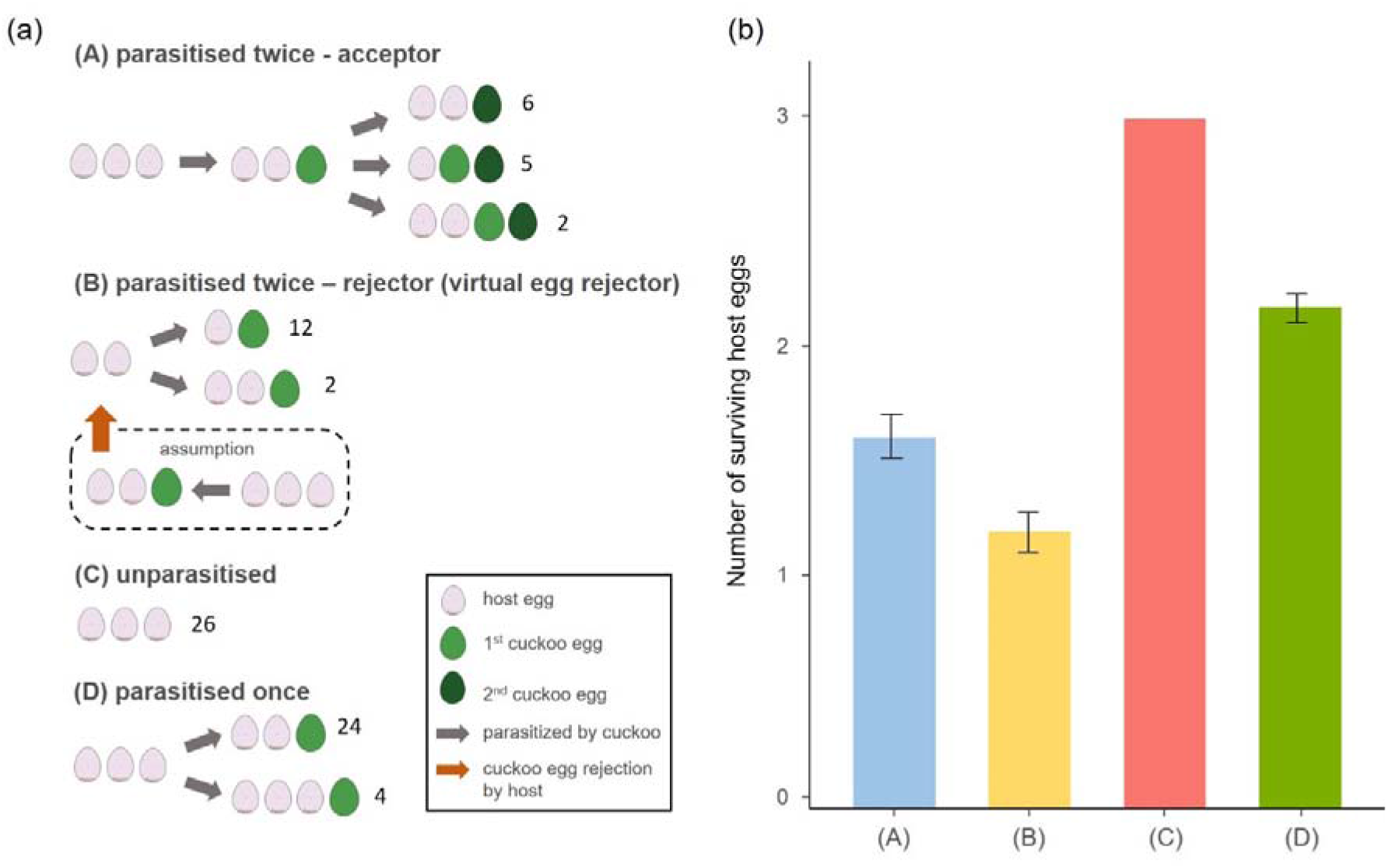
(a) An illustration of nest compositions (the numbers referring samples size) and (b) the number of host eggs remaining in nests at the end of incubation for nests of (A) multiply parasitised acceptor hosts, (B) virtual egg rejecter hosts, and (C) nests that were unparasitized or (D) singly parasitised. Host egg survival differed significantly between all groups.

Because gerygone clutches can naturally comprise either 2 or 3 eggs, it was not always possible to determine whether a cuckoo had removed the last gerygone egg (recently-laid 3^rd^ gerygone egg or no egg at all), as the parasitism event occurred on the same morning that a 3^rd^ gerygone egg would have been laid. That is, we could not confirm that the cuckoo encountered the treatment clutch of two gerygone eggs plus one cuckoo egg at the time of parasitism. We therefore excluded these nests from our analysis (n=11). For all groups, the number of surviving gerygone and cuckoo eggs was recorded at the end of the first week of incubation. To compare the number of surviving gerygone eggs in our four groups, we used a one-way ANOVA and pairwise comparisons.

We also recorded all cases of nest abandonment by gerygones that occurred during the egg stage of nesting. We used chi-square tests to compare the proportion of parasitized nests that were abandoned when at least one host egg remained, and when only cuckoo eggs (or one cuckoo egg) remained. To ensure equal opportunity for nest abandonment in all groups, we excluded nests from our dataset that did not survive until at least one week of incubation. All analyses were conducted using R ver. 3.5.3 (R Development Core Team, 2019) and the emmeans packages. Errors reported are the standard error of the mean.

## Results and Discussion

Among all large-billed gerygone nests, 66% (79 of 121) were parasitized by little bronze-cuckoos, and 34% (27 of 79) of these were parasitized with two (n=23) or more than two (n=4) cuckoo eggs. We confirm that for multiply parasitized nests of the large-billed gerygone, the presence of a first-laid cuckoo egg in the nest decreases the risk of host egg loss in a subsequent parasitism event, relative to a clutch in which that first cuckoo egg were rejected; multiply parasitized “acceptor” nests (those with two gerygone eggs and one cuckoo egg) retained 1.62 gerygone eggs (± 0.10, Fig. 1a (A) and 1b (A)) while “virtual egg rejecters” (those with two gerygone eggs only at the time of parasitism) retained 1.2 gerygone eggs (± 0.09, Fig. 1a (B) and 1b (B); ANOVA: df=3, *F*=97.73; *P*<0.000, All pairwise comparisons: *P*<0,05). This is because most cuckoos removed an egg at the time of laying (77 of 88 parasitism events) and around half of all second-to-arrive cuckoos removed a previously laid cuckoo egg, rather than a gerygone egg (n=9 of 18 multiple parasitism events). Singly parasitized nests retained 2.18 gerygone eggs (± 0.07, Fig. 1-b), and unparasitized nests had close to 100% egg survival.

In gerygones, the combination of high parasitism rate and small clutch size means that acceptance of cuckoo eggs poses another risk: that the gerygones are left tending a clutch comprising only cuckoo eggs. Interestingly, gerygones did abandon some parasitized nests (11%, 9 of 79) and they were more likely to abandon parasitized nests that contained only cuckoo egg/s (35%, 8 of 23) than those that retained at least one host egg (1%, 1 of 56; Pearson’s Chi-squared test with Yates’ continuity correction: df =1, χ^2^ = 14.64, *P*<0.000). However, gerygones never removed host or cuckoo eggs from their nests (unparasitized; n=42, parasitized; n=79). Nor did they ever abandon unparasitised nests containing host eggs (n=42). The trigger for nest abandonment appeared to be the absence of host eggs (rather than presence of the cuckoo egg), because gerygones were significantly more likely to abandon nests containing cuckoo eggs, but no host eggs, than nests containing both cuckoo eggs and host eggs. These results suggest that when parasitism rates are extremely high, it is beneficial for hosts to persevere with clutches containing at least one host egg, because any new breeding attempt is also likely to be parasitized. Hosts are also selected under such conditions though to recognize and reject nests in which no host eggs remain. Nest abandonment in large-billed gerygones thus appears to be a response to brood parasitism, and shows that large-billed gerygones exhibit plasticity in their egg stage defences, utilizing cuckoo egg acceptance in most cases (and benefiting from a clutch dilution effect), when there is some chance of successfully fledging their own young, but switching to nest abandonment when there is no possibility of rearing their own chicks.

Theoretical models propose that only egg acceptors will evolve chick discrimination (Britton et al., 2007; Grim, 2006; Planqué et al., 2002). At our study site, a parasitized gerygone nest has a one in three chance of being parasitized again. This high risk of egg loss from second-to-arrive cuckoos is coupled with the high cost of accepting cuckoo nestlings, whose eviction behaviour removes any remaining host brood. Thus, in gerygones and other systems with highly virulent parasites, multiple parasitism should promote chick rejection as the optimal host defense. In conclusion, our results demonstrate that multiple parasitism drives a facultative response to cuckoo eggs depending on clutch composition, and supports the argument that a clutch dilution effect has promoted the evolution of parasite nestling rejection (Sato et al., 2010).

## Acknowledgements

We are grateful to Virginia Abernathy, Garam Kim, Benjamin Ewing, Adélie Krellenstein, Owen Goodyear for their field assistance. We also acknowledge Cairns Regional Council for access to the Botanic Gardens and other council property. We kindly thank Brian Venables all his help with the little bronze-cuckoo project over several years

## Notes

### Competing Interest Statement

The authors have declared no competing interest.

